# *ACTIVA*: realistic single-cell RNA-seq generation with automatic cell-type identification using introspective variational autoencoders

**DOI:** 10.1101/2021.01.28.428725

**Authors:** A. Ali Heydari, Oscar A. Davalos, Lihong Zhao, Katrina K. Hoyer, Suzanne S. Sindi

## Abstract

**Motivation:** Single-cell RNA sequencing (scRNAseq) technologies allow for measurements of gene expression at a single-cell resolution. This provides researchers with a tremendous advantage for detecting heterogeneity, delineating cellular maps, or identifying rare subpopulations. However, a critical complication remains the low number of single-cell observations due to limitations by the rarity of a subpopulation, tissue degradation, or cost. This absence of sufficient data may cause inaccuracy or irreproducibility of downstream analysis. In this work, we present *ACTIVA* (Automated Cell-Type-informed Introspective Variational Autoencoder): a novel framework for generating realistic synthetic data using a single-stream adversarial variational autoencoder conditioned with cell-type information. Within a single framework, ACTIVA can generate data representative of the entire population, or specific subpopulations on demand, as opposed to two separate models (such as scGAN and cscGAN). Data generation and augmentation with ACTIVA can enhance scRNAseq pipelines and analysis, such as benchmarking new algorithms, studying the accuracy of classifiers, and detecting marker genes. ACTIVA will facilitate analysis of smaller datasets, potentially reducing the number of patients and animals necessary in initial studies.

**Results:** We train and evaluate models on multiple public scRNAseq datasets. In comparison to GAN-based models (scGAN and cscGAN), we demonstrate that ACTIVA generates cells that are more realistic and harder for classifiers to identify as synthetic, which also have better pair-wise correlations between genes. We show that data augmentation with ACTIVA significantly improves the classification of rare subtypes (more than 45% improvement compared to not augmenting and 4% better than cscGAN) all while reducing training time by an order of magnitude in comparison to both models.

**Availability of data and code:** Links to raw, pre- and post-processed data, source code and tutorials are available at https://github.com/SindiLab.

**Supplementary information:** Supplementary material can be found as a separate file with the same pre-print submission.

## Introduction

Traditional sequencing methods are limited to measuring the average signal in a group of cells, which potentially mask heterogeneity and rare populations [Tang *et al.* (2019)]. Single-Cell RNA sequencing (scRNAseq) technologies allow for the amplification and extraction of small RNA quantities, which enable sequencing at a single-cell level [Tang *et al.* (2009)]. The single-cell resolution thus enhances our understanding of complex biological systems. For example, in the immune system scRNAseq has been used to discover new immune cell populations, targets and relationships, which have been used to propose new treatments [Tang *et al.* (2019)].

While the number of tools for analyzing scRNAseq data increases, one limiting factor remains: low number of cells, potentially related to financial, ethical, or patient availability [Marouf *et al.* (2020)]. Large well-funded projects have generated the Human Cell Atlas [Regev *et al.* (2017)] and the Mouse Cell Atlas [Han *et al.* (2018)] which characterized cell populations in organs and tissues in their respective species. Although a tremendous amount of scRNAseq is available from these projects, they are limited to a broad overview of the cell populations in these tissues and organs. The Atlases overlook sub-populations of these cells which tend to be smaller, rarer, and important players in normal and dysregulated states. As Button *et al.* (2013) note, small numbers of observations reduce the reproducibility and robustness of experimental results. This is especially important for benchmarking new tools for scRNAseq data, as the number of features (genes) in each cell often exceeds the number of samples.

Given limitations on scRNAseq data availability and the importance of adequate sample sizes, *in-silico* data generation and augmentation offers a fast, reliable, and cheap solution. Synthetic data augmentation is a standard practice in fields of machine learning such as text and image classification [Shorten and Khoshgoftaar (2019)]. Traditional data augmentation techniques, geometric transformations or noise injection, are being replaced by more recently developed generative models, variational autoencoder (VAE) [Kingma and Welling (2013)] and Generative Adversarial Networks (GANs) [Goodfellow *et al.* (2014)], for augmenting complex biological datasets. However, GANs and VAEs remain less explored for data augmentation in genomics and transcriptomics. (We provide an overview of GANs and VAEs in Section 2.) Recently, Marouf *et al.* (2020) introduced GAN-based models for scRNAseq generation and augmentation (called single-cell GAN [scGAN] and conditional scGAN [cscGAN]), and demonstrated that they outperform other state-of-the-art models. While scGAN augments the entire population by creating “holistic” cells, cscGAN is conditioned to generate cells from specific subpopulations.

In this work, we extend and generalize their approach by employing an introspective VAE for data augmentation. The motivation and application of our generative model is closely related to Marouf et al.’s, with a focus on improving training time, stability and generation quality using only one framework. We compare our proposed model, ACTIVA, with scGAN and cscGAN and show how it can be leveraged to augment rare populations, improving classification and downstream analysis. In contrast to these previously published GANs, our novel cell-type conditioned introspective VAE model allows us to generate either “holistic” or specific cellular subpopulations in a single framework.

In Section 2 we provide an overview of GANs (both generally and in the context of scRNAseq data) and VAEs. In Section 3 we detail ACTIVA, our proposed conditional introspective VAE. In Section 4 we describe our training data and associated pre/post processing steps. In Section 5 we compare ACTIVA with competing methods - scGAN and cscGAN. We demonstrate that augmenting rare cell populations with ACTIVA improves classification over GANs while providing a more computationally tractable framework, mirroring both scGAN and cscGAN, in a single model. In comparison with scGAN and cscGAN, ACTIVA’s generates cells are harder for classifiers to identify as synthetic (i.e. having Areas Under the Curve closer to 0.5), with better pair-wise correlation between genes and that allow for improved classification of rare subtypes (more than 4% improvement over cscGAN) all while reducing run-time by an order of magnitude in comparison to both models. Finally, in Section 6, we review our approach, findings and limitations.

## Background

### 2.1 Generative Adversarial Networks

GANs [Goodfellow *et al.* (2014)] are capable of generating realistic synthetic data, and have been successfully applied to a wide range of machine learning tasks in computer vision [Dziugaite *et al.* (2015); Vondrick *et al.* (2016); Zhu *et al.* (2016)], natural language processing [Fedus *et al.* (2018); Yang *et al.* (2017b)], time series synthesis [Engel *et al.* (2019); Esteban *et al.* (2017)], and bioinformatics [Liu *et al.* (2019); Marouf *et al.* (2020)]. GANs consist of a generator network (*G*) and a discriminator network (*D*) that train adversarially, which enables them to produce high-quality fake samples. During training, *D* learns the difference between real and synthetic samples, while *G* produces fake data to “fool” *D*. More specifically, *G* produces a distribution of generated samples *P_g_*, given an input *z ~ P_z_*, with *P_z_* being a random noise distribution. The objective of GANs is to learn *P_g_*, ideally finding a close approximation to the real data distribution *P_r_*, so that *P_g_ ≈ P_r_*. To learn the approximation to *P_g_*, GANs play a “min-max game” of

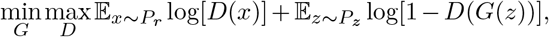

where both players (*G* and *D*) attempt to maximize their own payoff. This adversarial training of *G* and *D* is critical in GANs’ ability to generate realistic samples. Compared to other generative models, GANs’ main advantages are (i) the ability to produce any type of probability density, (ii) no prior assumptions for training the generator network, and (iii) no restrictions on the size of the latent space.

Despite these advantages, GANs are notoriously hard to train since it is highly non-trivial for *G* and *D* to achieve Nash equilibrium [Wang *et al.* (2019)]. Another disadvantage of GANs are vanishing gradients where an optimal *D* cannot provide enough information for *G* to learn and make progress. That is, if *D* learns the distinction between real and generated data too well, then *G* will fail to train, as shown by Arjovsky and Bottou (2017). Another issue with GANs is “mode collapse”, when *G* generates only a small set of outputs that can trick *D*. More explicitly, this occurs when *G* has learned to map several noise vectors *z* to the same output that *D* classifies as real data. In this scenario, *G* is over-optimized, and the generated samples lack diversity. Quantifying how much GANs have learned the distribution of real data is often complicated, consisting of measuring the dissimilarity between *P_g_* and *P_r_* when *P_r_* is not known or assumed. Therefore, common ways of evaluating GANs involve direct evaluation of the output [Larsen *et al.* (2016)], which can be arduous.

Although some variations of GANs have been proposed to alleviate vanishing gradients and mode collapse (e.g. Wasserstein-GANs (WGANs) [Arjovsky *et al.* (2017)] and Unrolled-GANs [Metz *et al.* (2016)]), the convergence of GANs still remains a major problem. During the training progression, the feedback of *D* to *G* becomes meaningless, and if GANs continue to train past this point, the quality of the synthetic samples can be affected and ultimately collapse. It is also important to note that commonly used GANs cannot be trained as single-stream networks, and a necessary step is to define a training schedule for *G* and *D* separately, adding another layer of complexity. Although all deep learning models are sensitive to hyperparameter choices, Lucic *et al.* (2018) show all experimented GANs (including WGANs) are much more sensitive to these choices than VAEs. This can be a drawback in using GANs for scRNAseq generation since the hyperparameters may need to be re-tuned for every new dataset.

### 2.2 Single Cell GANs

scGAN and cscGAN are the state-of-the-art deep learning models for generating and augmenting scRNAseq data. Marouf et al. train scGAN to generate single-cell data from all populations and cscGAN to produce cluster-specific samples, with the underlying model in both being a WGAN. For scGAN, the objective is to minimize Wasserstein distance between real cells distribution, *P_r_*, and generated data, *P_g_*:

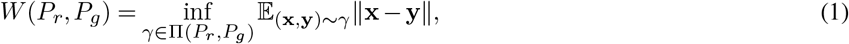

where **x** and **y** denote random variables, II(*P_r_, P_g_*) is the set of all joint probability distributions *γ*(**x**, **y**) with marginals *P_r_* and *P_g_*. Intuitively, Wasserstein distance is the cost of optimally transporting “masses” from **x** to **y** such that *P_r_* is transformed to *P_g_* [Arjovsky *et al.* (2017)]. However, since the infimum in Eq. (1) is highly intractable, Arjovsky et al. use Kantorovich-Rubinstein duality to find an equivalent formulation of Wasserstein distance with better properties:

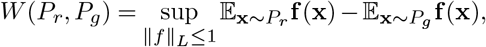

where the set of 1-Lipschitz functions is denoted by ||*f*||_*L*_ ≤ 1, with the solution being a universal approximator (potentially a fully connected neural network) to approximate *f*. This function is approximated by *D*, which we denote as *f_d_*. Similarly, we denote the function approximated by the generator as *f_g_*. Using these notations, we arrive at the adversarial objective function of WGANs (used in scGAN):

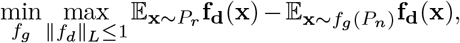

where *P_n_* denotes a multivariate noise distribution.

As mentioned earlier, training a WGAN instead of a traditional GAN can alleviate the vanishing gradient issue. However, the majority of GANs’ training instabilities can still be seen in WGANs, making WGANs less flexible and transferable between different datasets or domain-specific tasks. cscGAN uses a projection-based conditioning [Miyato and Koyama (2018)] which adds an inner product of class labels (cell types) at the discriminator’s output. Based on instruction given by the authors in the implementation, scGAN and cscGAN must be trained separately; however, our model learns to generate specific cell populations (cell-types) or collective cell clusters with one training.

### 2.3 Variational Autoencoders

VAEs [Kingma and Welling (2013, 2019)] are generative models that jointly learn deep latent-variable and inference models, i.e. they are comprised of a generative model and an inference model. VAEs are autoencoders that use variational inference to reconstruct the original data, giving the ability to generate new (or random) data that is “similar” to those already in a dataset *x*. VAEs assume that observed data and latent representation are jointly distributed as *P_θ_*(*x, z*)= *P_θ_*(*x|z*)*P*(*z*). In deep learning, the log-likelihood *P_θ_*(*x|z*) is defined as a network *Gen*(·) with parameters *θ*, and a noise model with *ϵ* ~ *P*(*ϵ*) where *x*|*z* = *Gen_θ_*(·)+*ϵ*. Due to the intractability of maximizing the expected log-likelihood of observed data over *θ*, 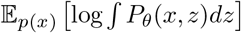, the goal is to instead maximize the evidence lower bound (ELBO):

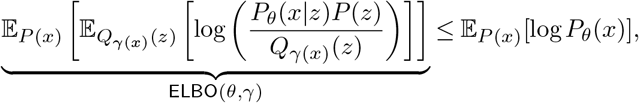

where *Q*_*γ*(*x*)_(*z*) is an auxiliary variational distribution (with parameters *γ*(*x*)) that tries to approximate the true posterior *P_θ_*(*z|x*).

Note that when *Q*_*γ*(*x*)_(*z*)= *P_θ_*(*z|x*), the lower bound approaches the expected log-likelihood, therefore aiming to infer *P_θ_*(*z*|*x*). We find the variational parameters *γ* for new inputs *x* using an inference network *Enc*(·) with parameters *ϕ*, such that *Enc_ϕ_*(*x*)= *γ*(*x*). As shown by Zhao *et al.* (2017), this maximization problem can be written as

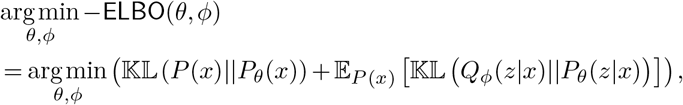

where *Q_ϕ_*(*z*|*x*) denotes the variational distributions *Enc_ϕ_*(*x*) and 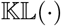 denotes the Kullback–Leibler (KL) divergence. This yields to the objective function

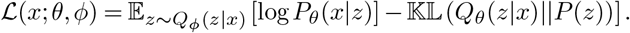

ELBO is the underlying objective function for this work, modified by Huang *et al.* (2018) to allow VAEs to train in an adversarial manner while maintaining key mathematical properties, as described in Section 3.

The main issue with VAEs arises when the training procedure falls into the trivial local optimum of the ELBO objective; that is, when the variational posterior and the true posterior closely match the prior (or collapse to the prior). This phenomenon often causes issues with data generation since the generative model ignores a subset of latent variables that may have meaningful latent features for inputs [He *et al.* (2019)]. This arises because of the 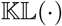 term in the VAE’s objective function [Lucas *et al.* (2019)], i.e., when the divergence between the prior and the posterior is close to zero. Studies that aim to alleviate posterior collapse can be broadly categorized as (i) solutions aiming to weaken the generative model [Semeniuta *et al.* (2017); Yang *et al.* (2017a)], (ii) modifications to the training objective [Tolstikhin *et al.* (2018); Zhao *et al.* (2018)], and (iii) modifications to the training procedure [He *et al.* (2019); Heydari *et al.* (2019)]. In our experiments, we did not encounter posterior collapse. However, in our package, we provide an option for modifying objective function weights adaptively using SoftAdapt [Heydari *et al.* (2019)].

VAEs have also been criticized for generating samples that are “blurry” (adhering to an average of the data points), as opposed to sharp samples that GANs produce because of adversarial training. This issue has often been addressed by defining an adversarial training between the encoder and the decoder, as done in introspective VAEs (IntroVAEs) [Huang *et al.* (2018)], which we use in our framework. IntroVAEs are single-stream generative models that self-evaluate the quality of the generated images. They have been used mostly in computer vision, which have performed comparably to their GAN counterparts, in applications such as synthetic image generation [Huang *et al.* (2018)] and single-image super-resolution [Heydari and Mehmood (2020)]. We describe the formulations of IntroVAEs and the individual components of our model in Section 3, and provide a visual representation in Fig. 1.

**Fig. 1.**
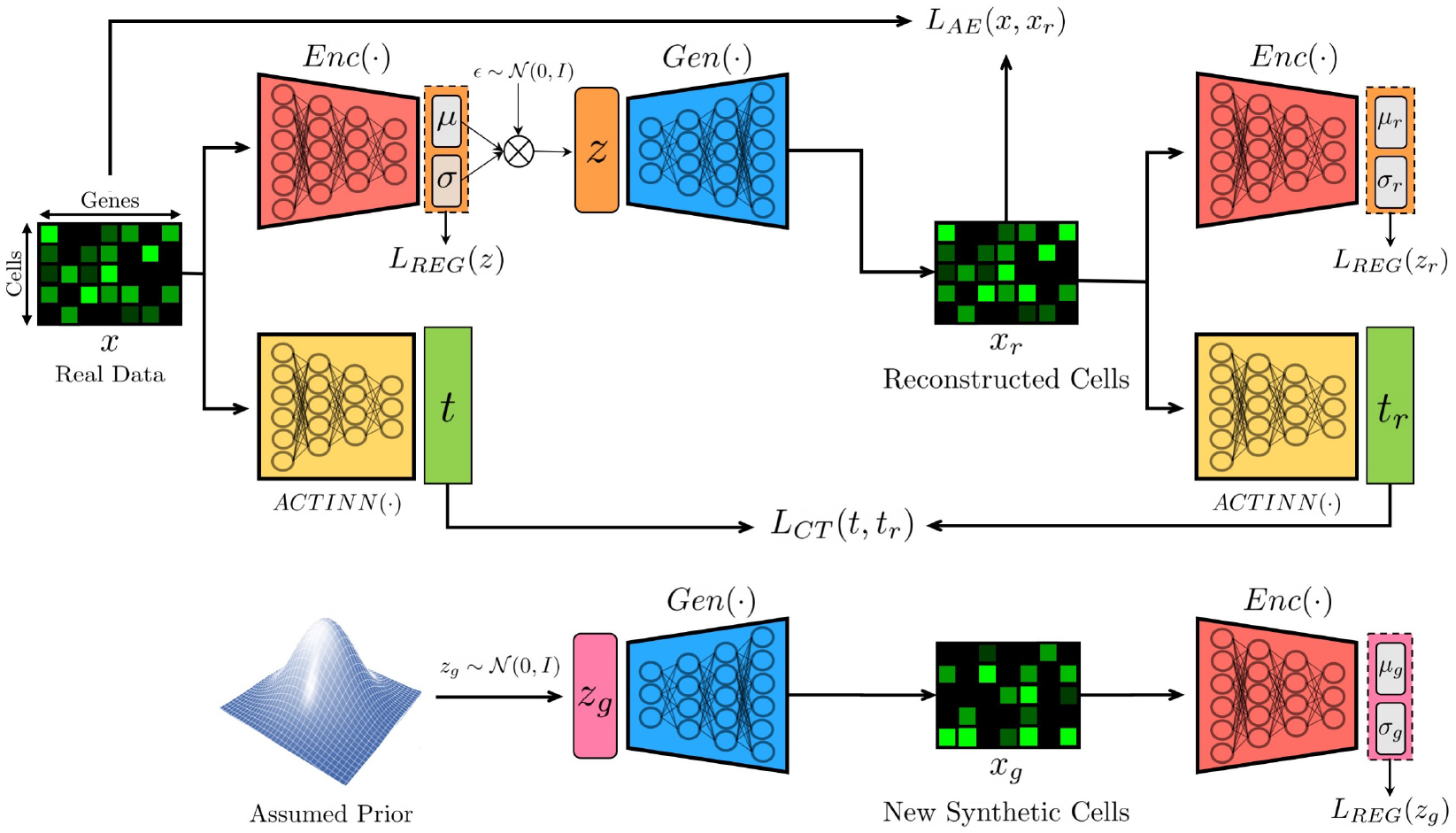
Overview of our deep generative model, ACTIVA, and its training flow. Our model consists of three networks: (i) an encoder that also acts as a discriminator (denoted by *Enc*), (ii) a decoder which is used as the generative network for producing new synthetic data (denoted by *Gen*), and (iii) an automated cell-type classification network (denoted by ACTINN as in [Ma and Pellegrini (2019)]). ACTIVA is a single-stream adversarial (Introspective) Conditional VAE without the need for a training schedule (unlike most GANs). Due to this, our model has the training stability and efficiency of VAEs while producing realistic samples comparably to GANs. We describe each component of our model and the objective functions in Section 3.

VAEs’ natural ability to produce both a generative and an inference model presents them as an ideal candidate for generation and augmentation of omics data. In this work, we demonstrate the ability of our deep VAE-based model for producing realistic in-silico scRNAseq data. Our model, ACTIVA, performs comparably to the state-of-the-art GAN models, scGAN and cscGAN, and trains significantly faster and maintains stability. Moreover, ACTIVA learns to generate specific cell-types and holistic population data in one training (unlike scGAN and cscGAN that train separately). On the same datasets and in the same environment, our model trains at least 6 times faster than scGAN. Moreover, ACTIVA can produce 100K samples in less than 2 seconds on a single NVIDIA Tesla V100, and 87 seconds on a common research laptop. ACTIVA provides researchers with a fast, flexible, and reliable deep learning model for augmenting and enlarging existing datasets, improving downstream analyses robustness and reproducibility.

## Methods and Approach

Our proposed model, ACTIVA, consists of three main networks, with a self-evaluating VAE as its core and a cell-type classifier as its conditioner. In this section, we formulate the objective functions of our model and describe the training procedure.

### 3.1 Encoder Network

The ACTIVA encoder network, *Enc*, serves two purposes: (i) mapping (encoding) scRNAseq data into an approximate posterior to match the assumed prior, and (ii) acting as a discriminator, judging the quality of the generated samples against training data. Therefore, *Enc*’s objective function is designed to train as an adversary of the generator network, resulting in realistic data generation. To approximate the prior distribution, KL divergence is used as a regularization term (denoted as *L_REG_*) which regularizes the encoder by forcing the approximate posterior, *Q_ϕ_*(*z*|*x*), to match the prior, *P* (*z*) (following notation from Section 1). In this work, we assume a center isotropic multivariate Gaussian prior, but it is important to note that studies have considered negative binomial or zero-inflated binomials as the prior for scRNAseq data [Anders and Huber (2010); Robinson and Smyth (2007)].

Given our assumption of the prior, the posterior probability is 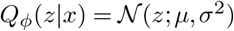, where *μ* and *σ* are the mean and standard deviation, respectively, computed from the outputs of *Enc*. As in traditional VAEs, *z* is sampled from 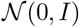 which will be used as an input to the generator network (decoder in VAEs). Due to the stochasticity of *z*, gradient-based backpropagation becomes difficult, but using the reparameterization trick in Kingma and Welling (2013) makes this operation tractable. That is, define *z* = *μ* + *σ* ⊙ *ϵ* with 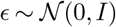 which passes the stochasticity of *z* onto *ϵ*. Now given *N* cells and a latent vector in a *D*-dimensional space (i.e. *z* ∈ ℝ^*D*^), we can compute the KL regularization:

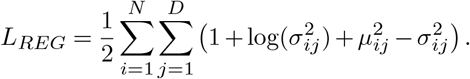

Similar to traditional VAEs, the encoder network aims at minimizing the difference between reconstructed and training cells (real data). We denote this reconstruction loss as *L_AE_*. As in Huang *et al.* (2018), our reconstruction objective is to minimize the mean squared error between training cells *x* ∈ ℝ^*M*^ (*M* being the number of genes/features) and reconstructed cells *x_r_* ∈ ℝ^*M*^:

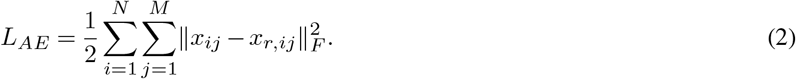

As the last part of the network, we introduce a cell-type loss component that encourages *x_r_* to have the same cell type as *x*. That is, given a classifying network *C*, we want to ensure that the identified type of reconstructed sample *C*(*x_r_*)= *t_r_* is the same as real cell *C*(*x*)= *t*. For this, we introduce *L_CT_*, shown in Eq. (5). We describe the explicit formulation and the classifying network in Section 3.3. During the development of ACTIVA, Zheng *et al.* (2020) introduced a similar conditioning loss to the IntroVAE framework for image synthesis, which provided significant improvements in generating new data (images).

Given our model’s objectives, the loss function for *Enc*, 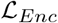, must encode training data and self-evaluate newly generated cells from the generator network *Gen*:

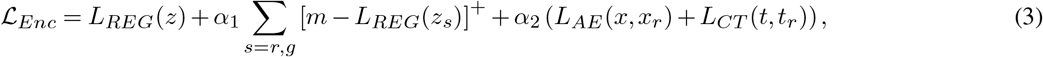

where subscripts *r* and *g* denote reconstructed and generated cells from *Gen*, respectively. Note that reconstructed cells *x_r_* correspond directly to training data *x*, but generated cells *x_g_* are newly produced cells. In Eq. (3), [·]^+^ = max(0, ·), and *m* ∈ ℝ^+^ determines our network’s adversarial training, as described in Section 3.4.

### 3.2 Generator Network

Generator network, *Gen*, aims at learning two tasks. First, *Gen* must learn a mapping of encoded training data, *z* ∈ ℝ^*D*^, from the posterior, *Q_ϕ_*(*z*|*x*), back to the original feature space, ℝ^*M*^. In the ideal mapping, reconstructed samples *x_r_* would match training data *x* perfectly. To encourage learning of this objective, we minimize the mean squared error between *x* and *x_r_*, as shown in Eq. (2), and force cell-types of reconstructed samples to match with original cells, as shown in Eq. (5). The second task of *Gen* is to generate realistic new samples from a random noise vector *z_n_* ∈ ℝ^*D*^ ~ *P*(*z*) (sampled from the prior *P* (*z*)) that “fool” the encoder network *Enc*. That is, after producing new synthetic samples *x_g_*, we calculate *Enc*(*x_g_*)= *z_g_* to judge the quality of generated cells. Given these two objectives, the generator’s objective function is defined as

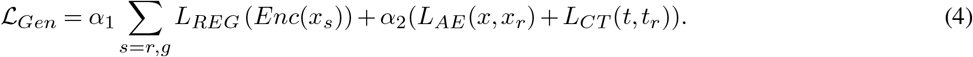

### 3.3 Automated Cell Type Conditioning

Minimizing *L_AE_* alone does not encourage producing diverse samples from all populations explicitly, and for this reason, we introduce a cell-type matching objective. The goal of this objective is to encourage the generator to generate cells that are classified as the same type as the input data. More explicitly, the loss component *L_CT_* will penalize the network if reconstructed cell-types are different from the training data. Given a trained classifier *C*(·), we can express this objective as

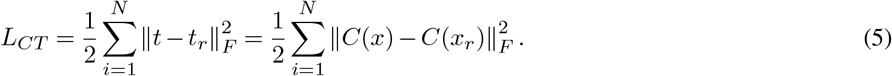

For ACTIVA’s conditioning, we use an automated cell-type identification introduced by Ma and Pellegrini (2019). This network, called ACTINN, uses all genes to capture features, and focuses on the signals associated with cell variance. We chose ACTINN because of its accurate classification and efficiency in training compared to other existing models [Abdelaal *et al.* (2019)]; we provide an overview of ACTINN in the Supplementary Material. Our model is also flexible to use any classifier as a conditioner, as long as an explicit loss could be computed between the predicted labels and the true labels^1^. With ACTINN as the classifier, *t* and *t_r_* are logits (output layer) for *x* and *x_r_* respectively. Our implementation of ACTINN is available as a stand-alone package at https://github.com/SindiLab/ACTINN-PyTorch.

### 3.4 Adversarial Training

The generator produces two types of synthetic cells: reconstructed cells *x_r_* from *x* and newly generated cells *x_g_* from a noise vector. While both the *Enc* and *Gen* attempt to minimize *L_AE_* and *L_CT_*, the encoder tries to minimize *L_REG_*(*z*) and maximize *L_REG_*(*z_r,g_*) to be greater than or equal to *m*. However, the generator tries to minimize *L_REG_*(*z_r,g_*) in order to minimize its objective function. This is the min-max game played by *Enc* and *Gen*. Note that choosing *m* is an important step for the network’s adversarial training; we describe the strategies for the choice of *m* in the Supplementary Material.

### 3.5 Network Architecture

Our network architecture is shown in Fig. 2, with the input *x* ∈ ℝ^*M*^. Latent vectors of our model live in a 128-dimensional space, but for the sake of generality, we assume *z* ∈ ℝ^*D*^. In the encoder and generator networks, Adam [Kingma and Ba (2015)] optimizer is used with a learning rate *lr* = 0.0002, and moving averages decay rates *β*_1_ = 0.9, *β*_2_ = 0.999. Gradients are calculated on mini-batches of size 128 with the adversarial constant *m* = 110. We describe each component in more detail in the Supplementary Material.

**Fig. 2.**
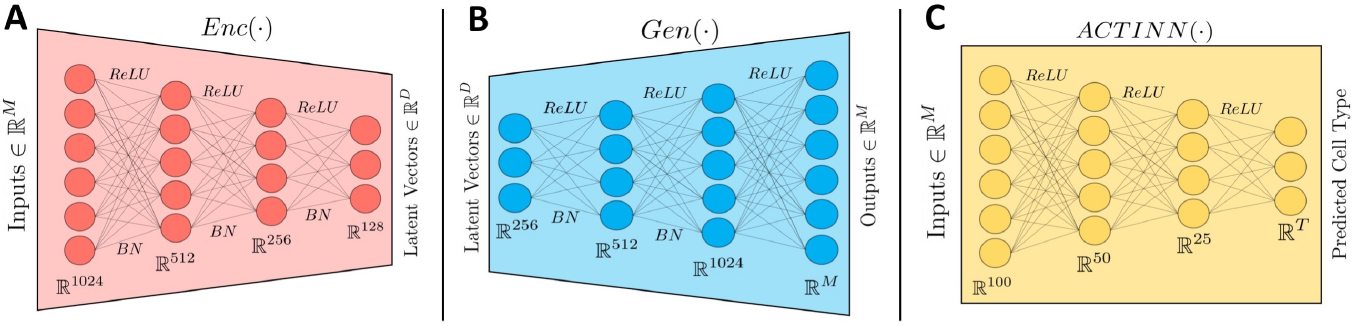
Overview of ACTIVA networks. Here we present **A**, the encoder network, **B** the generator network, and **C**, the cell-type classifier. *ReLU* denotes Rectified Linear Units [Nair and Hinton (2010)], and *BN* denotes batch normalization operation [Ioffe and Szegedy (2015)].

### 3.6 Training and Inference Procedure

Unlike GANs, our model does not require a training schedule. The cell-type classifier in ACTIVA can be pre-trained or trained simultaneously with *Enc* and *Gen*. To start the adversarial training and pick an appropriate *m*, we first train the model as a VAE for 10 epochs while training ACTINN in parallel (these options are readily available and adjustable in our package). After the initial warm-up, we train the IntroVAE component for 600 epochs with *α*_1_ =1 and *α*_2_ = 0.5 (from Eq. (3)-(4)). We also provide an option for dynamic weight-balancing using SoftAdapt that would be useful in the case of a posterior collapse (which did not occur in our experiments).

For inference, we input a random noise tensor sampled from a multi-variate Gaussian to the trained generator. For cell-specific generation, outputs are automatically filtered through the trained classifier to produce the desired sub-populations on demand. Tutorials and notebooks on training and inference are available via link provided in abstract.

## Datasets and Data Processing

### 4.1 Pre-Processing

We use the pipeline provided by Marouf et al. to pre-process the data. First, we removed genes that were expressed in less than 3 cells and cells that expressed less than 10 genes. Next, cells were normalized by total unique molecular identifiers (UMI) counts and scaled to 20000 reads per cell. Then, we selected a “test set” (approximately 10% of each dataset). Testing samples were randomly chosen considering cell ratios in each cluster (“balanced split”). Links to raw and pre-processed datasets are available via the link provided in abstract.

### 4.2 Datasets

#### 68K PBMC

To compare our results with the current state-of-the-art deep learning model, scGAN/cscGAN, we trained and evaluated our model on a dataset containing 68579 peripheral blood mononuclear cells (PBMCs) from a healthy donor (68K PBMC) [Zheng *et al.* (2017)]. 68K PBMC is an ideal dataset for evaluating generative models due to the distinct cell populations, data complexity, and size [Marouf *et al.* (2020)]. After pre-processing, the data contained 17789 genes. We then performed a balanced split on this data, which resulted in 6991 testing and 61588 training cells.

#### Brain Small

In addition to 68K PBMC, we used a randomly-selected subset of a larger dataset called Brain Large (both by 10x Genomics). Brain small contains 20,000 random samples (out of approximately 1.3 million cells) from the cortex, hippocampus, and the subventricular zone of two embryonic day 18 mice. Compared to 68K PBMC, this dataset has fewer cells, and it varies in complexity and organism. The full dataset and and its subset, Brain Small, are available on 10X Genomics portal. After performing the pre-processing steps, the data contained 17970 genes, which we then split (via “balanced split”) to 1997 test cells and 18003 training cells.

#### 4.3 Post-Processing

After generating a count matrix with a generative model (e.g. ACTIVA or scGAN), we add the gene names (from the real data) and save as a Scanpy/Seurat object. We then use Seurat to identify 3000 highly variable genes through the use of variance-stabilization transformation (VST) [Hafemeister and Satija (2019)], which applies a negative binomial regression to identify outlier genes. The shared highly variable genes are then used for integration [Stuart *et al.* (2019)], which allows for biological feature overlap between different datasets in order to perform the downstream analyses presented in this work. Next, we perform a gene-level scaling, i.e. centering the mean of each feature to zero and scaling by the standard deviation. The feature space is then reduced to 50 principal components, followed by Uniform Manifold Approximation and Projection (UMAP) [McInnes *et al.* (2018)] and t-distributed Stochastic Neighbor Embedding (t-SNE) [van der Maaten and Hinton (2008)]. As noted by Marouf *et al.* (2020), analysis with lower-dimensional representations have two main advantages: (i) most biologically relevant information is captured while noise is reduced and (ii) statistically, it is more acceptable to use lower dimensional embeddings in classification tasks when samples and features are of the same order of magnitude, which is often the case with scRNAseq datasets (such as the ones we used). Lastly, we use Scater [McCarthy *et al.* (2017)] to visualize the datasets.

## Results

Assessing generative model quality is notoriously difficult and still remains an open research area [Lucic *et al.* (2018); Theis *et al.* (2016)]. Here, we apply some qualitative and quantitative metrics for evaluating synthetic scRNAseq, as used in Marouf *et al.* (2020). For qualitative metrics, we compare the manifold of generated and real cells using UMAP. For quantitative metrics, we train a classifier to distinguish between real and synthetic cells. To study ACTIVA’s performance, we compare our results to Marouf et al. alone since their models outperform other state-of-the-art generative models such as Splatter [Zappia *et al.* (2017)] and SUGAR [Lindenbaum *et al.* (2018)]. Training and inference time comparisons are shown in Supplementary Material. As we show, ACTIVA generates cells that better resemble the real data, and it outperforms competing methods on improving classification of rare cell populations with data augmentation. ACTIVA is one model that can be served as an alternative to both scGAN and cscGAN, and it trains much faster than both GAN-based models (and it only needs one training).

### 5.1 ACTIVA Trains Faster than GAN-Based Models

To measure the efficiency of ACTIVA in comparison to the state-of-the-art GAN-based models (scGAN and cscGAN), we trained all three models in the exact same computational environment on a single GPU for each dataset (we describe the hardware used in Computational Environment section). Note that since scGAN and cscGAN train separately, we repeated this process five times to account for any variability, and then computed the average training time and standard deviation. As shown in Table 1, ACTIVA trains orders of magnitude faster for both datasets (approximately 17 times faster on Brain Small and 6 times faster on 68K PBMC) and only needs one training to produce cells from all populations (scGAN’s aim) and specific cell populations (cscGAN’s purpose).

**Table 1.**
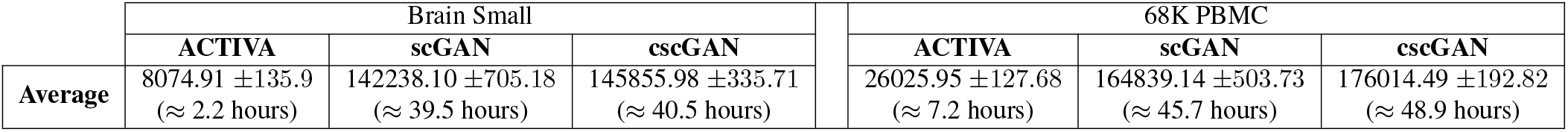
Average training time with sample standard deviation (in seconds) for ACTIVA, scGAN and cscGAN in the same computational environment. Here, we report the average of five training times for each model. ACTIVA (which has the capabilities of scGAN and cscGAN combined) trains much faster than both scGAN and cscGAN (approximately 17 times faster on Brain Small and 6 times faster on 68K PBMC, on average). Individual run-times for each iteration are provided in Supplementary Material.

|  | Brain Small |  |  | 68K PBMC |  |  |
| --- | --- | --- | --- | --- | --- | --- |
|  | ACTIVA | scGAN | cscGAN | ACTIVA | scGAN | cscGAN |
| Average | 8074.91 ± 135.9<br>(≈ 2.2 hours) | 142238.10 ± 705.18<br>(≈ 39.5 hours) | 145855.98 ± 335.71<br>(≈ 40.5 hours) | 26025.95 ± 127.68<br>(≈ 7.2 hours) | 164839.14 ± 503.73<br>(≈ 45.7 hours) | 176014.49 ± 192.82<br>(≈ 48.9 hours) |

### 5.2 ACTIVA Generates Realistic Cells

To qualitatively evaluate the generated cells, we analyzed the two-dimensional UMAP representation of the test set (real data) and in-silico generated cells (same size as the test set). We found that the distribution and clusters match closely between ACTIVA generated cells and real cells (Fig. 4A–4B and Fig. S7). We also analyzed t-SNE embeddings of the real cells and synthetic cells generated by ACTIVA, which showed similar results. These qualitative assessments demonstrated that ACTIVA learns the underlying manifold of real data, the main goal of generative models. A key feature of ACTIVA is the cell-type conditioning which encourages the network to produce cells from all clusters. This means that generating cells with ACTIVA results in *gaining cells within clusters rather than losing clusters*. Due to this design choice, ACTIVA can generate more cells from the rare populations than scGAN, as shown in Fig. 4A-4B. ACTIVA’s flexible framework allows for adjusting the strength of the cell-type conditioning (which is a parameter in our model) for the cases where the exact data representation is more desirable.

Next we quantitatively assessed the quality of the generated cells by training a random forest (RF) classifier (same as in Marouf *et al.* (2020)) to distinguish between real and generated cells. The goal here is to determine how “realistic” ACTIVA generated cells are compared to real cells. Ideally, the classifier will not differentiate between the synthetic and real cells, thus resulting in a receiver operating characteristic (ROC) curve that is the same as randomly guessing (0.5 area under the curve [AUC]). The RF classifier consists of 1000 trees with the Gini impurity criterion and the square root of the number of genes as the maximum number of features used. Maximum depth is set to either all leaves containing less than two samples or until all leaves are pure. We generated cells using ACTIVA and scGAN and performed a five-fold cross-validation on synthetic and real cells (test). ACTIVA performs better than scGAN with AUC scores closer to 0.5 for both datasets (Fig. 3). For Brain Small test set, the mean ACTIVA AUC is 0.62 ± 0.02 compared to scGAN’s 0.74 ± 0.01. For 68K PBMC, the mean AUC is 0.68 ± 0.01 for ACTIVA and 0.73 ± 0.01 for scGAN.

**Fig. 3.**
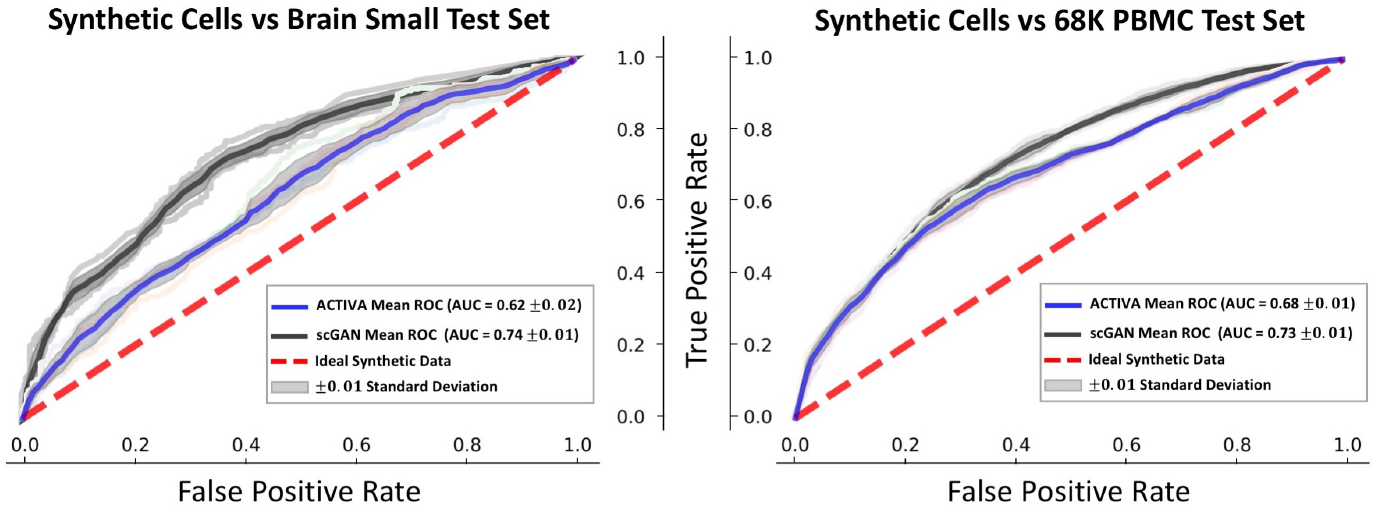
Classifying synthetic data (ACTIVA and scGAN) from real data (test set) for Brain Small (left plot) and 68K PBMC (right plot). The metrics are reported using a random forest classifier (detailed in Section 5.2) with 5 fold cross-validation (marked by pastel colors in each plot). An area under the curve (AUC) of 0.5 (chance) is the ideal scenario (red dash line), and an AUC closer to this value is better.

### 5.3 ACTIVA Generates Similar Gene Expression Profiles

In order to generate cells that represent all clusters, the marker gene distribution in the generated data should roughly match the gene distribution in real cells. We used UMAP representations of ACTIVA, scGAN, and the test set, and colored them based on the expression levels of marker genes. Fig. 4C–4D show examples of log-gene expression for marker genes from each datasets (additional examples in Supplementary Material). In our qualitative assessment, ACTIVA generated cells following the real gene expression closely. For a quantitative assessment, we calculated the Pearson correlation of top 5 differentially expressed genes from each cluster for both ACTIVA generated cells and real data. As shown in Fig. 5, the pairwise correlation of genes from the ACTIVA generated cells closely match those from the real data for both datasets. To quantify the overall gene-gene correlation for synthetic data, we define a metric to measure the discrepancy between the correlations of generated data and the real data (test set). Specifically, given a correlation matrix of generated samples, *G*, and a correlation matrix of the real data, *R*, we compute the 1-norm of the difference in correlations to measure the performance of the synthetic data, shown in Eq. (6), which we refer to as *Correlation Discrepancy* (CD) for simplicity:

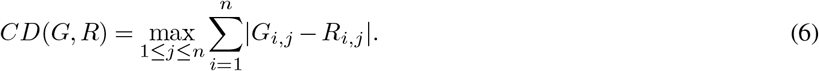

(In the ideal case, *CD*(*G, R*)= 0, therefore values closer to zero indicate better performance.) Using Eq. (6), we find that for 68K PBMC, ACTIVA has a CD score of 1.5816, as opposed to scGAN’s 2.2037, and for Brain Small ACTIVA outperforms scGAN with a CD score of 4.6852 compared to 5.5937. These values further quantify that generated cells from ACTIVA better preserve the gene-gene correlation present in the real data.

**Fig. 4.**
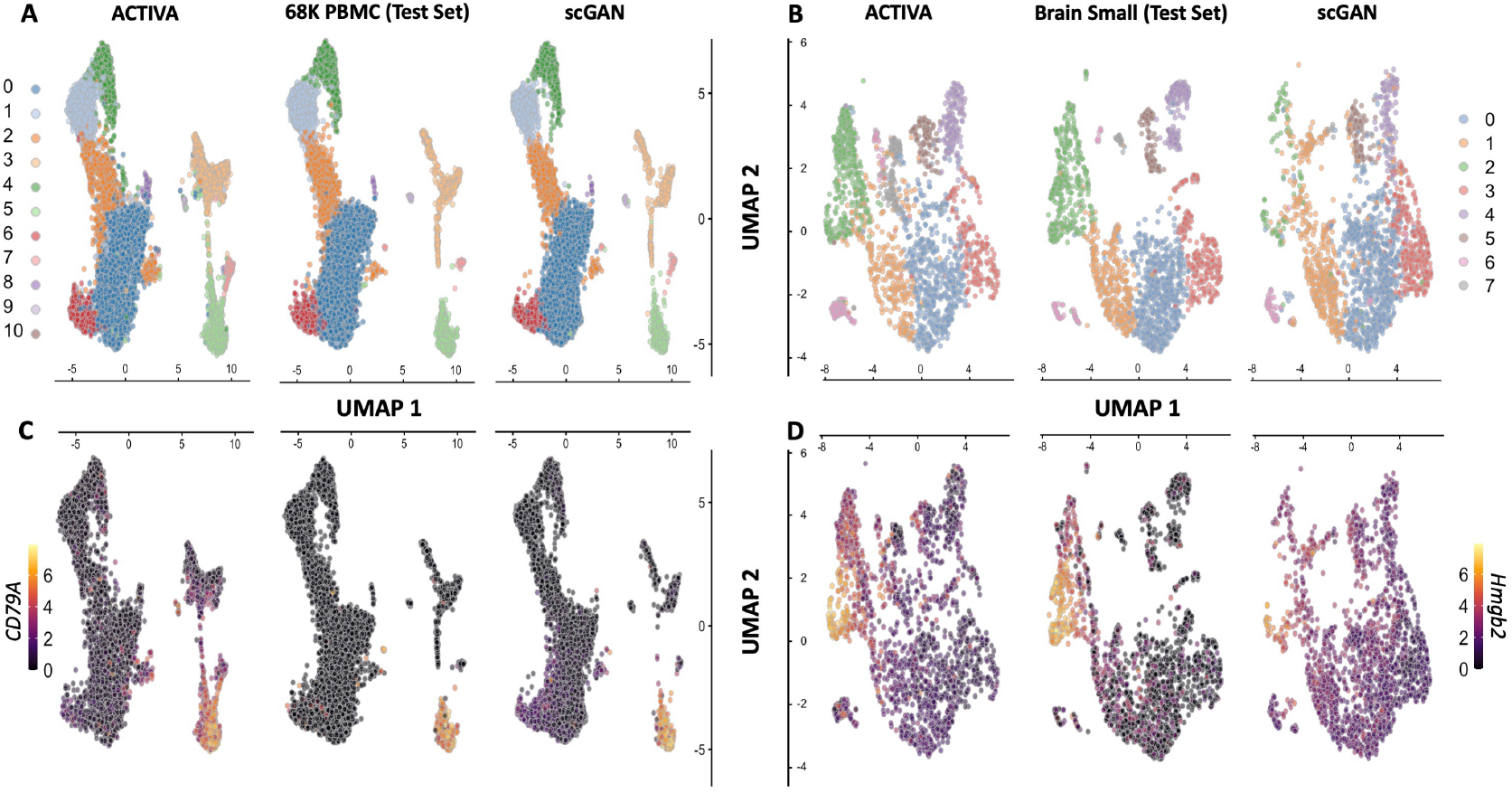
ACTIVA generates high-quality cells that resemble both the cluster and gene expressions present in the training data. Top row: UMAP plot of ACTIVA generated cells compared with test set and scGAN generated cells, colored by clusters for 68K PBMC (**A**) and Brain Small (**B**). Bottom row: same UMAP plots as top row, colored by selected marker gene expressions. (**C**) corresponds the log expression for CD79A marker gene (for 68K PBMC) and (**D**) illustrates the same for Hmgb2 (for Brain Small). ACTIVA’s cell-type conditioning encourages to generate more cells per cluster rather than lose clusters, meaning that ACTIVA will generate more cells from the rare populations (e.g. cluster 7 of PBMC and cluster 6 of Brain Small).

**Fig. 5.**
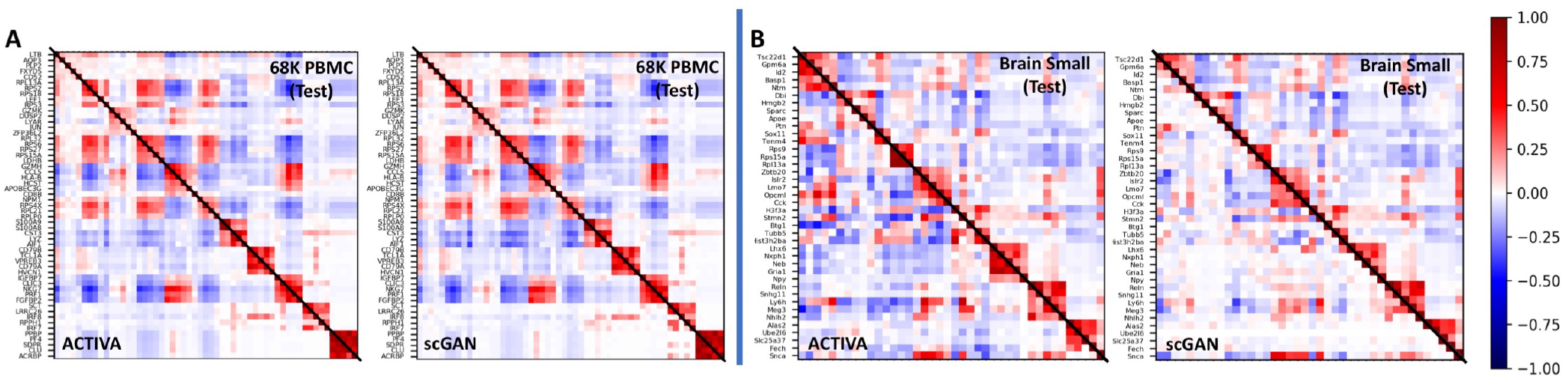
Correlation of top five differentially expressed genes in each cluster for 68K PBMC (A) and Brain Small (B). Lower triangular of matrices indicate correlation of generated data, and upper triangular show correlation of real data (same in both plots in panels A and B). For 68K PBMC (**A**), we investigated pairwise correlation for a total of 55 genes and for Brain Small (**B**), we calculated the Pearson correlation for 40 genes. In the ideal case, the correlation plots should be symmetric, and the Correlation Discrepancy (CD), the metric defined in Eq. (6), should be zero. The gene correlations in ACTIVA match the real data more closely than scGAN, as shown above and as computed through CD; ACTIVA has a CD score of 1.5816 and 4.6852 for 68K PBMC and Brain Small, respectively, compared to scGAN’s 2.2037 and 5.5937.

Additionally, we plotted marker gene distribution in all cells against real cells. Fig. S6 illustrates the distribution of five marker genes from cluster 1 (LTB, LDHB, RPL11, RPL32, RPL13) and cluster 2 (CCL5, NKG7, GZMA, CST7, CTSW). We also investigated known marker genes for specific cell populations, such as for B-cells in PBMC data, finding that ACTIVA generated cells expressed these markers (CD79A, CD19, and MS4A1) in the appropriate clusters (Fig. 4C and other figures not shown here) similar to real data. Following Marouf et al., we calculated the maximum mean discrepancy (MMD) between the real data distribution and the generated ones using ACTIVA and scGAN. Simply stated, MMD is a distance metric based on embedding probabilities in a reproducing kernel Hilbert space [Gretton *et al.* (2012)], and since MMD is a distance metric, a lower value of MMD between two distributions indicates the distribution are closer to one another. For consistency, we chose the same kernels as Marouf et al. and calculated MMD on the first 50 principal components. As shown in Table 2, ACTIVA had a lower MMD score than scGAN, demonstrating an improvement in the quality of the generated cells compared to scGAN.

**Table 2.** MMD values for ACTIVA, scGAN and positive control (training set) compared to the test set. In this metric, ACTIVA outperforms scGAN for both datasets, since it has a lower MMD score.

|  | Brain Small |  |  | 68K PBMC |  |  |
| --- | --- | --- | --- | --- | --- | --- |
|  | Training Set | ACTIVA | scGAN | Training Set | ACTIVA | scGAN |
| MMD | 0.0619 | 0.7715 | 0.9686 | 0.0539 | 0.7952 | 0.9047 |

Based on qualitative and quantitative evaluations of our model, we conclude that ACTIVA has learned the underlying marker gene distribution of real data, as desired. However, we suspect that assuming a different prior in model formulation (e.g., Zero-Inflated Negative Binomial) could further improve our model’s learning of real data.

### 5.4 ACTIVA Generates Specific Cell-Types on Demand

Since we minimize a cell-type identification loss in the training objective, ACTIVA is encouraged to produce cells that are classified correctly. Therefore, the accuracy of the generated cell-types depends on the classifier selected. In Tables S4 and S5, we show that ACTIVA’s classifier accurately distinguishes rare cell-types, achieving an F1 score of 0.89 when trained with only 1% sub-population in the training cells. ACTIVA generates specific cell-types from the manifold it has learned, which then filters through the identifier network to produce specific sub-populations. To quantify the quality of the generated samples, we trained an RF classifier (as in Section 5.2) to distinguish between generated and real sub-populations in the data. Fig. 6 illustrates ACTIVA’s performance against cscGAN for the Brain Small dataset, which shows ACTIVA achieving better AUC scores. Similar results were obtained for 68K PBMC sub-populations, although the AUC gap between cscGAN and ACTIVA were narrower.

**Fig. 6.**
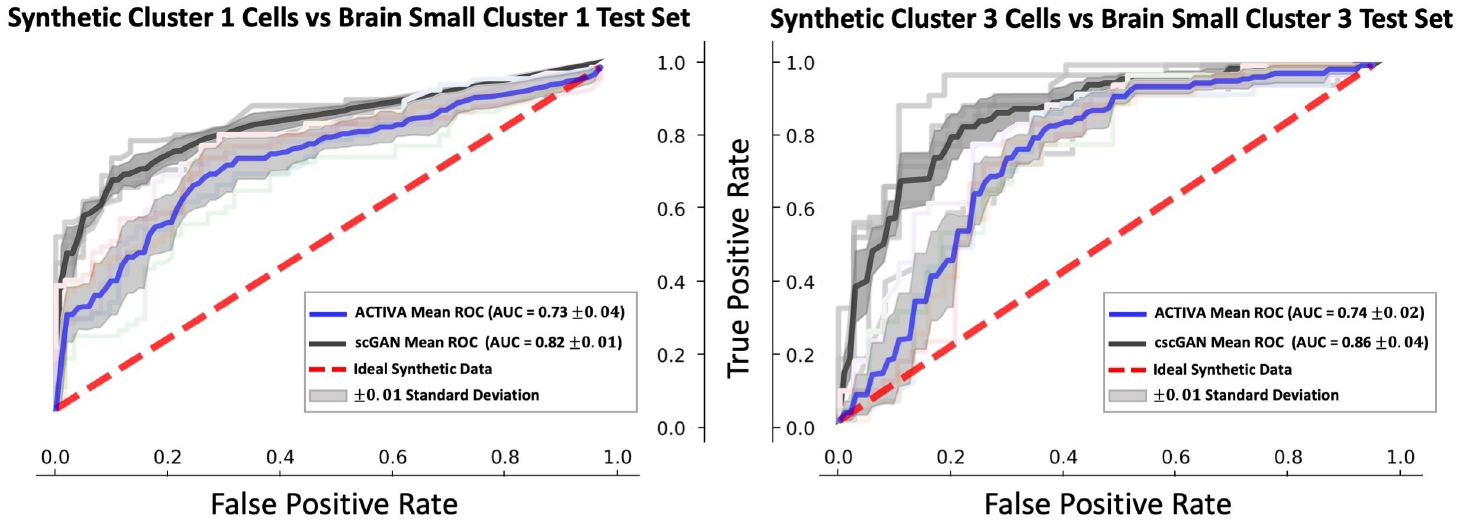
RF classifier distinguishes two real sub-populations from synthetic data for Brain Small. ACTIVA out-performs cscGAN in producing realistic samples on this dataset, since the ROC curve is closer to chance (red dashed line).

### 5.5 ACTIVA Improves Classification of Rare Cells

A main goal of designing generative models is to augment sparse datasets with additional data that can improve downstream analyses. Given the performance of our model and conditioner, we hypothesized that classifying rare cells in a dataset can be improved through augmentation with ACTIVA, i.e. using synthetic rare cells alongside real data. We next directly compared against cscGAN to demonstrate the feasibility of augmenting rare population to improve classification. We utilized the data-augmentation experiment presented by Marouf *et al.* (2020). That is, we chose the cells in cluster 2 of 68K PBMC, and downsampled those cells to 10%, 5%, 1% and 0.5% of the actual cluster size, while keeping the populations fixed. The workflow of the downsampling and exact sizes is shown in Fig. S4 (Supplementary Material). We then trained ACTIVA on the downsampled subsets, and generated 1500 synthetic cluster 2 cells to augment the data (Marouf et al. generated 5000 cells). After that, we used an RF classifier to identify cluster 2 cells versus all other cells. This classification was done on (i) downsampled cells without augmentation and (ii) downsampled cells with ACTIVA augmentation. F1 scores are measured on a held-out test set (10% of the total real cluster 2 cells), shown in Fig. 7. The classifier is identical to the one described in Section 5.2 with the addition of accounting cluster-size imbalance, as it was done by Marouf et al, since RF classifiers are sensitive to unbalanced classes [Zadrozny *et al.* (2003)]. Most notably, our results show an improvement of 0.4526 in F1 score (from 0.4736 to 0.9262) when augmenting 0.5% of real cells, and an improvement of 0.2568 (from 0.6829 to 0.9397) on the 1% dataset. ACTIVA also outperforms augmentation with cscGAN for the rarest case, since cscGAN achieves an F1 score of 0.8774 as opposed to ACTIVA’s 0.9262. These results indicate a promising and powerful application of ACTIVA in rare cell-type identification and classification.

**Fig. 7.**
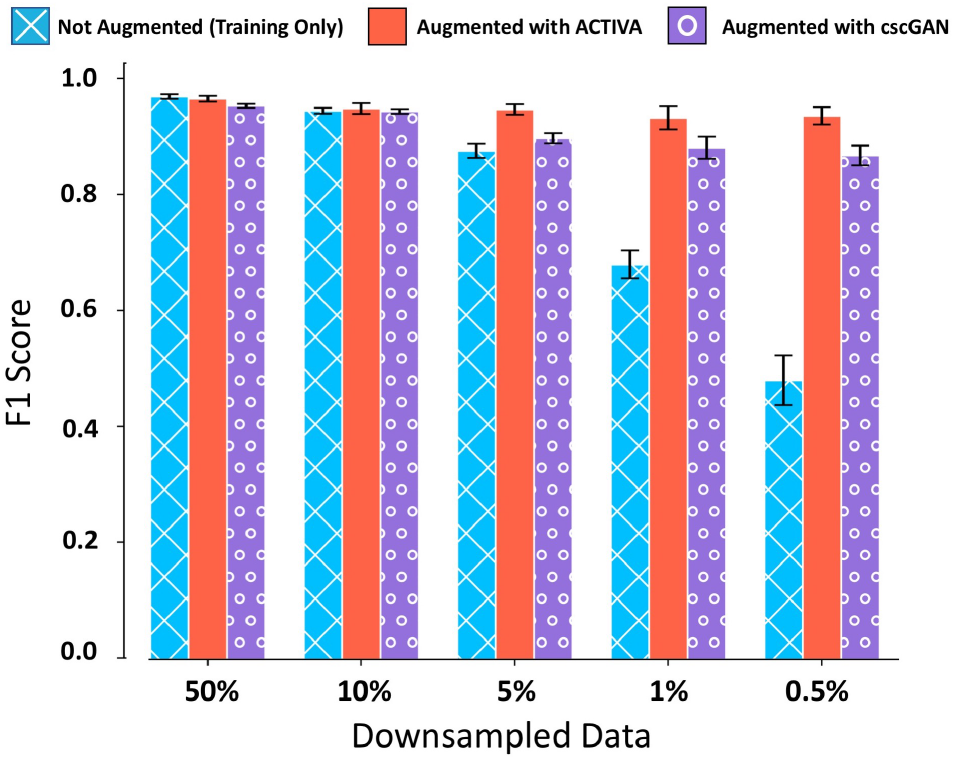
Augmentation with ACTIVA improves classification of rare populations. Mean F1 scores of RF classifier for training data (with no augmentation), shown in blue, and training data with augmentation, shown in red. Error bars indicate the range for five different random seeds for sub-sampling cluster 2 cells.

## Conclusions and Discussion

In this manuscript, we propose a deep generative model for generating realistic scRNAseq data. Our model, ACTIVA, consists of an automatic cell-type identification network, coupled with IntroVAE that aims to learn the distribution of the original data and the existing sub-populations. Due to the architectural choices and single-stream training, ACTIVA trains orders of magnitude faster than the state-of-the-art GAN-based model, and produces samples that are comparably in quality. ACTIVA can be easily trained on different datasets to either enlarge the entire dataset (generate samples from all clusters) or augment specific rare populations.

ACTIVA can generate hundreds of thousands of cells in only a few seconds (on a GPU), which enables benchmarking of new scRNAseq tools’ accuracy and scalability. We showed that, for these datasets, using ACTIVA for augmenting rare populations improves downstream classification by more than 40% in the rarest case of real cells used (0.5% of the training samples). We believe that ACTIVA learns the underlying higher dimensional manifold of the scRNAseq data, even where there are few cells available. The deliberate architectural choices of ACTIVA provide insights as to why this learning occurs. As Marouf et al. also noted, the fully connected layers of our three networks share information learned from all other populations. In fact, the only cluster-specific parameters are the ones learned in the batch normalization layer. This is also shown with the accuracy of the conditioner network when trained on rare populations. However, if the type-identifying network does not classify sub-populations accurately, this can directly affect the performance of the generator and inference model due to the conditioning. We keep this fact in mind and therefore allow for the flexibility of adding any classifier to our existing architecture.

Lopez *et al.* (2018) demonstrate that the latent manifold of VAEs can also be useful for analyses such as clustering or denoising. Deep investigation of the learned manifold of ACTIVA can further improve the interpretability of our model, or yield new research questions to explore. We also hypothesize that assuming a different prior such as a Zero Inflated Negative Binomial or a Poisson distribution could further improve the quality of generated data. Our experiments show that ACTIVA learns to generate high-quality samples on complex datasets from different species. ACTIVA potentially reduces the need for human and animal sample sizes and sequencing depth in studies, saving costs and time, and improving robustness of scRNAseq research with smaller datasets. Furthermore, ACTIVA would benefit studies where large or diverse patient sample sizes are not available, such rare and emerging disease.

## Computational Environment

Development and testing were done on Accelerated Computing EC2 instances (`p3.2xlarge and p3.8xlarge`) of Amazon Web Services. All requirements and dependencies are automatically installed by our package and are listed in a requirements file, but for the sake of completeness, they are as follows: Python v3.7.6, PyTorch v1.5.1, NumPy v1.18.5, SciPy v1.4.1, Pandas v1.2.0, Scanpy v1.6.0, AnnData v0.7.5, and Scikit-learn v0.24.0. For data pre and post processing, we used LoomPy v3.0.6, SeuratDisk v0.0.0.9013, Seurat v3.2.3, and R v4.0.3. The scGAN package was run in a Docker container. Reported training times for ACTIVA/scGAN were averages of 5 times on a single NVIDIA-Tesla V100 GPU. Inference times were averages of 5 measurements on (i) V100 GPU (GPU time) and (ii) 2.3 GHz Quad-Core Intel Core i7 (on a 2020 MacBook Pro).

## Acknowledgements

We acknowledge Stefan Bonn and Pierre Machart for their support with scGAN, & Maia Powell and Alina Gataullina for helpful feedback.

## Footnotes

1 This is true for most classifiers.

## References

Abdelaal, T., Michielsen, L., Cats, D., Hoogduin, D., Mei, H., Reinders, M. J. T., and Mahfouz, A. (2019). A comparison of automatic cell identification methods for single-cell RNA sequencing data. Genome Biology, 20(1), 194.

Anders, S. and Huber, W. (2010). Differential expression analysis for sequence count data. Genome Biology, 11(10), R106.

Arjovsky, M. and Bottou, L. (2017). Towards principled methods for training generative adversarial networks. arXiv:1701.04862.

Arjovsky, M., Chintala, S., and Bottou, L. (2017). Wasserstein generative adversarial networks. volume 70 of Proceedings of Machine Learning Research, pages 214–223, International Convention Centre, Sydney, Australia. PMLR.

Button, K. S., Ioannidis, J. P., Mokrysz, C., Nosek, B. A., Flint, J., Robinson, E. S., and Munafò, M. R. (2013). Power failure: why small sample size undermines the reliability of neuroscience. Nature reviews neuroscience, 14(5), 365–376.

Dziugaite, G. K., Roy, D. M., and Ghahramani, Z. (2015). Training generative neural networks via maximum mean discrepancy optimization. arXiv preprint arXiv:1505.03906.

Engel, J., Agrawal, K. K., Chen, S., Gulrajani, I., Donahue, C., and Roberts, A. (2019). Gansynth: Adversarial neural audio synthesis. arXiv preprint arXiv:1902.08710.

Esteban, C., Hyland, S. L., and Rätsch, G. (2017). Real-valued (medical) time series generation with recurrent conditional GANs. arXiv preprint arXiv:1706.02633.

Fedus, W., Goodfellow, I., and Dai, A. M. (2018). MaskGAN: better text generation via filling in the_. arXiv preprint arXiv:1801.07736.

Goodfellow, I., Pouget-Abadie, J., Mirza, M., Xu, B., Warde-Farley, D., Ozair, S., Courville, A., and Bengio, Y. (2014). Generative adversarial nets. In Advances in neural information processing systems, pages 2672–2680.

Gretton, A., Borgwardt, K. M., Rasch, M. J., Schölkopf, B., and Smola, A. (2012). A kernel two-sample test. Journal of Machine Learning Research, 13(25), 723–773.

Hafemeister, C. and Satija, R. (2019). Normalization and variance stabilization of single-cell RNA-seq data using regularized negative binomial regression. Genome Biology, 20(1), 296.

Han, X., Wang, R., Zhou, Y., Fei, L., Sun, H., Lai, S., Saadatpour, A., Zhou, Z., Chen, H., Ye, F., Huang, D., Xu, Y., Huang, W., Jiang, M., Jiang, X., Mao, J., Chen, Y., Lu, C., Xie, J., Fang, Q., Wang, Y., Yue, R., Li, T., Huang, H., Orkin, S. H., Yuan, G.-C., Chen, M., and Guo, G. (2018). Mapping the mouse cell atlas by microwell-seq. Cell, 172(5), 1091–1107.

He, J., Spokoyny, D., Neubig, G., and Berg-Kirkpatrick, T. (2019). Lagging inference networks and posterior collapse in variational autoencoders. In International Conference on Learning Representations.

Heydari, A. A. and Mehmood, A. (2020). Srvae: super resolution using variational autoencoders. In Pattern Recognition and Tracking XXXI, volume 11400, page 114000U. International Society for Optics and Photonics.

Heydari, A. A., Thompson, C. A., and Mehmood, A. (2019). Softadapt: Techniques for adaptive loss weighting of neural networks with multi-part loss functions. CoRR, abs/1912.12355.

Huang, H., Li, Z., He, R., Sun, Z., and Tan, T. (2018). IntroVAE: Introspective variational autoencoders for photographic image synthesis. In S. Bengio, H. Wallach, H. Larochelle, K. Grauman, N. Cesa-Bianchi, and R. Garnett, editors, Advances in Neural Information Processing Systems 31, pages 52–63. Curran Associates, Inc.

Ioffe, S. and Szegedy, C. (2015). Batch normalization: Accelerating deep network training by reducing internal covariate shift. In F. Bach and D. Blei, editors, Proceedings of the 32nd International Conference on Machine Learning, volume 37 of Proceedings of Machine Learning Research, pages 448–456, Lille, France. PMLR.

Kingma, D. P. and Ba, J. (2015). Adam: A method for stochastic optimization. In Y. Bengio and Y. LeCun, editors, 3rd International Conference on Learning Representations, ICLR 2015, San Diego, CA, USA, May 7-9, 2015, Conference Track Proceedings.

Kingma, D. P. and Welling, M. (2013). Auto-encoding variational Bayes. CoRR, abs/1312.6114.

Kingma, D. P. and Welling, M. (2019). An introduction to variational autoencoders. Foundations and Trends^®^ in Machine Learning, 12(4), 307–392.

Larsen, A. B. L., Sønderby, S. K., Larochelle, H., and Winther, O. (2016). Autoencoding beyond pixels using a learned similarity metric. volume 48 of Proceedings of Machine Learning Research, pages 1558–1566, New York, New York, USA. PMLR.

Lindenbaum, O., Stanley, J., Wolf, G., and Krishnaswamy, S. (2018). Geometry based data generation. In S. Bengio, H. Wallach, H. Larochelle, K. Grauman, N. Cesa-Bianchi, and R. Garnett, editors, Advances in Neural Information Processing Systems, volume 31, pages 1400–1411. Curran Associates, Inc.

Liu, Q., Lv, H., and Jiang, R. (2019). hicGAN infers super resolution Hi-C data with generative adversarial networks. Bioinformatics, 35(14), i99–i107.

Lopez, R., Regier, J., Cole, M. B., Jordan, M. I., and Yosef, N. (2018). Deep generative modeling for single-cell transcriptomics. Nature Methods, 15(12), 1053–1058.

Lucas, J., Tucker, G., Grosse, R. B., and Norouzi, M. (2019). Don’t blame the ELBO! A linear VAE perspective on posterior collapse. In H. Wallach, H. Larochelle, A. Beygelzimer, F. d’Alché-Buc, E. Fox, and R. Garnett, editors, Advances in Neural Information Processing Systems, volume 32, pages 9408–9418. Curran Associates, Inc.

Lucic, M., Kurach, K., Michalski, M., Bousquet, O., and Gelly, S. (2018). Are GANs created equal? A large-scale study. In Proceedings of the 32nd International Conference on Neural Information Processing Systems, NIPS’18, page 698–707, Red Hook, NY, USA. Curran Associates Inc.

Ma, F. and Pellegrini, M. (2019). ACTINN: automated identification of cell types in single cell RNA sequencing. Bioinformatics, 36(2), 533–538.

Marouf, M., Machart, P., Bansal, V., Kilian, C., Magruder, D. S., Krebs, C. F., and Bonn, S. (2020). Realistic in silico generation and augmentation of single-cell RNA-seq data using generative adversarial networks. Nature Communications, 11(1), 166.

McCarthy, D. J., Campbell, K. R., Lun, A. T. L., and Wills, Q. F. (2017). Scater: pre-processing, quality control, normalization and visualization of single-cell RNA-seq data in R. Bioinformatics, 33(8), 1179–1186.

McInnes, L., Healy, J., and Melville, J. (2018). UMAP: Uniform manifold approximation and projection for dimension reduction.

Metz, L., Poole, B., Pfau, D., and Sohl-Dickstein, J. (2016). Unrolled generative adversarial networks. CoRR, abs/1611.02163.

Miyato, T. and Koyama, M. (2018). cGANs with projection discriminator. In International Conference on Learning Representations.

Nair, V. and Hinton, G. E. (2010). Rectified linear units improve restricted boltzmann machines. In Proceedings of the 27th International Conference on International Conference on Machine Learning, ICML’10, page 807–814, Madison, WI, USA. Omnipress.

Regev, A., Teichmann, S. A., Lander, E. S., Amit, I., Benoist, C., Birney, E., Bodenmiller, B., Campbell, P., Carninci, P., Clatworthy, M., Clevers, H., Deplancke, B., Dunham, I., Eberwine, J., Eils, R., Enard, W., Farmer, A., Fugger, L., Göttgens, B., Hacohen, N., Haniffa, M., Hemberg, M., Kim, S., Klenerman, P., Kriegstein, A., Lein, E., Linnarsson, S., Lundberg, E., Lundeberg, J., Majumder, P., Marioni, J. C., Merad, M., Mhlanga, M., Nawijn, M., Netea, M., Nolan, G., Pe’er, D., Phillipakis, A., Ponting, C. P., Quake, S., Reik, W., Rozenblatt-Rosen, O., Sanes, J., Satija, R., Schumacher, T. N., Shalek, A., Shapiro, E., Sharma, P., Shin, J. W., Stegle, O., Stratton, M., Stubbington, M. J. T., Theis, F. J., Uhlen, M., van Oudenaarden, A., Wagner, A., Watt, F., Weissman, J., Wold, B., Xavier, R., and Yosef, N. (2017). The human cell atlas. Elife, 6.

Robinson, M. D. and Smyth, G. K. (2007). Moderated statistical tests for assessing differences in tag abundance. Bioinformatics, 23(21), 2881–2887.

Semeniuta, S., Severyn, A., and Barth, E. (2017). A hybrid convolutional variational autoencoder for text generation. In Proceedings of the 2017 Conference on Empirical Methods in Natural Language Processing, pages 627–637, Copenhagen, Denmark. Association for Computational Linguistics.

Shorten, C. and Khoshgoftaar, T. M. (2019). A survey on image data augmentation for deep learning. Journal of Big Data, 6(1), 60.

Stuart, T., Butler, A., Hoffman, P., Hafemeister, C., Papalexi, E., Mauck, W. M., Hao, Y., Stoeckius, M., Smibert, P., and Satija, R. (2019). Comprehensive integration of single-cell data. Cell, 177(7), 1888–1902.e21.

Tang, F., Barbacioru, C., Wang, Y., Nordman, E., Lee, C., Xu, N., Wang, X., Bodeau, J., Tuch, B. B., Siddiqui, A., et al. (2009). mRNA-seq whole-transcriptome analysis of a single cell. Nature methods, 6(5), 377–382.

Tang, X., Huang, Y., Lei, J., Luo, H., and Zhu, X. (2019). The single-cell sequencing: new developments and medical applications. Cell & Bioscience, 9(1), 53.

Theis, L., van den Oord, A., and Bethge, M. (2016). A note on the evaluation of generative models. In International Conference on Learning Representations.

Tolstikhin, I., Bousquet, O., Gelly, S., and Schoelkopf, B. (2018). Wasserstein auto-encoders. In International Conference on Learning Representations.

van der Maaten, L. and Hinton, G. (2008). Visualizing data using t-SNE. Journal of Machine Learning Research, 9(86), 2579–2605.

Vondrick, C., Pirsiavash, H., and Torralba, A. (2016). Generating videos with scene dynamics. In Advances In Neural Information Processing Systems, pages 613–621.

Wang, Z., She, Q., and Ward, T. E. (2019). Generative adversarial networks in computer vision: A survey and taxonomy. arXiv preprint arXiv:1906.01529.

Yang, Z., Hu, Z., Salakhutdinov, R., and Berg-Kirkpatrick, T. (2017a). Improved variational autoencoders for text modeling using dilated convolutions. In International conference on machine learning, pages 3881–3890. PMLR.

Yang, Z., Hu, J., Salakhutdinov, R., and Cohen, W. (2017b). Semi-supervised QA with generative domain-adaptive nets. In Proceedings of the 55th Annual Meeting of the Association for Computational Linguistics (Volume 1: Long Papers), pages 1040–1050, Vancouver, Canada. Association for Computational Linguistics.

Zadrozny, B., Langford, J., and Abe, N. (2003). Cost-sensitive learning by cost-proportionate example weighting. In Third IEEE International Conference on Data Mining, pages 435–442.

Zappia, L., Phipson, B., and Oshlack, A. (2017). Splatter: simulation of single-cell RNA sequencing data. Genome Biology, 18(1), 174.

Zhao, J., Mathieu, M., and LeCun, Y. (2016). Energy-based generative adversarial network. *arXiv*:1609.03126.

Zhao, S., Song, J., and Ermon, S. (2017). Towards deeper understanding of variational autoencoding models. *arXiv*:1702.08658.

Zhao, S., Song, J., and Ermon, S. (2018). InfoVAE: Information maximizing variational autoencoders. *arXiv*:1706.02262.

Zheng, G. X. Y., Terry, J. M., Belgrader, P., Ryvkin, P., Bent, Z. W., Wilson, R., Ziraldo, S. B., Wheeler, T. D., McDermott, G. P., Zhu, J., Gregory, M. T., Shuga, J., Montesclaros, L., Underwood, J. G., Masquelier, D. A., Nishimura, S. Y., Schnall-Levin, M., Wyatt, P. W., Hindson, C. M., Bharadwaj, R., Wong, A., Ness, K. D., Beppu, L. W., Deeg, H. J., McFarland, C., Loeb, K. R., Valente, W. J., Ericson, N. G., Stevens, E. A., Radich, J. P., Mikkelsen, T. S., Hindson, B. J., and Bielas, J. H. (2017). Massively parallel digital transcriptional profiling of single cells. Nature Communications, 8(1), 14049.

Zheng, K., Cheng, Y., Kang, X., Yao, H., and Tian, T. (2020). Conditional introspective variational autoencoder for image synthesis. IEEE Access, 8, 153905–153913.

Zhu, J.-Y., Krähenbühl, P., Shechtman, E., and Efros, A. A. (2016). Generative visual manipulation on the natural image manifold. In European Conference on Computer Vision, pages 597–613. Springer.

