## Supplementary Material for "*ACTIVA*: realistic single-cell RNA-seq generation with automatic cell-type identification using introspective variational autoencoders"

*Heydari et al.*

### Complete Computational Environment

Development and testing were done on Accelerated Computing EC2 instances (`p3.2xlarge` and `p3.8xlarge`) of Amazon Web Services. All requirements and dependencies are automatically installed by our package and are listed in a requirements file, but for the sake of completeness, they are as follows: Python v3.7.6, PyTorch v1.5.1, NumPy v1.18.5, SciPy v1.4.1, Pandas v1.2.0, Scanpy v1.6.0, AnnData v0.7.5, and Scikit-learn v0.24.0. For data pre- and post-processing, we used LoomPy v3.0.6, SeuratDisk v0.0.0.9013, Seurat v3.2.3, scater v1.16.2, and R v4.0.3. The scGAN package was run in a Docker container (using the provided dockerfile at https://github.com/imsb-uke/scGAN/tree/master/dockerfile). Reported training times for ACTIVA/scGAN were averages of 5 times on a single NVIDIA-Tesla V100 GPU (Table S1). Inference times were averages of 5 measurements on (i) V100 GPU (GPU time) and (ii) 2.3 GHz Quad-Core Intel Core i7 (on a 2020 MacBook Pro).

### GPU Training Times

**Table S1.** Training time (in seconds) on 68K PBMC for ACTIVA and scGAN on 1 NVIDIA Tesla V100 GPU. We trained each model 5 times under the same conditions to find an average training time. We have not included cscGAN times since training that model took longer than scGAN. We can see that ACTIVA trains much faster (6.3 times faster on average). scGAN and cscGAN were run in a Docker container (by Marouf et al.) using the Dockerfile located at https://github.com/imsb-uke/scGAN/tree/master/dockerfile).

| Iteration | ACTIVA | scGAN | cscGAN |
|---|---|---|---|
| 1 | 25968.2447 | 164225.5618 | 175978.2588 |
| 2 | 26218.7306 | 165308.7142 | 176041.7922 |
| 3 | 25935.2738 | 164391.8113 | 176318.0701 |
| 4 | 26091.8337 | 165281.5137 | 175941.7097 |
| 5 | 25915.6733 | 164988.1371 | 175792.6410 |
| Average | 26025.95 ($\approx$7.2 hours) | 164839.14 ($\approx$45.7 hours) | 176014.49 ($\approx$48.8 hours) |

**Table S2.** Training time (in seconds) on Brain Small for ACTIVA and scGAN on 1 NVIDIA Tesla V100 GPU. We trained each model 5 times under the same conditions to find an average training time. ACTIVA trains approximately 17 times faster than scGAN. scGAN and cscGAN were run in a Docker container (using the Dockerfile located at https://github.com/imsb-uke/scGAN/tree/master/dockerfile).

| Iteration | ACTIVA | scGAN | cscGAN |
|---|---|---|---|
| 1 | 8277.4867 | 142371.9793 | 145980.3154 |
| 2 | 7922.1005 | 141103.8196 | 145952.2580 |
| 3 | 8107.3591 | 143008.7401 | 145261.1851 |
| 4 | 7983.4031 | 142532.3804 | 146076.8253 |
| 5 | 8084.2473 | 142173.5900 | 146009.3626 |
| Average | 8074.91 ($\approx$2.2 hours) | 142238.10 ($\approx$39.5 hours) | 145855.98 ($\approx$40.5 hours) |

### Choosing Adversarial Constant $m$

Ensuring the numerical balance between the KL divergence regularization of real and fake samples is crucial to ACTIVA's sample quality. Therefore an $m$ that is extremely large or small could result affect how realistic the generated cells are. The adversarial training in IntroVAE is similar to Energy-based GANs [Zhao *et al.* (2016)]. The following are two strategies we followed for choosing an appropriate value of $m$, based on Zhao *et al.* (2016):

1. An effective strategy is to train the model as VAE for $n$ epochs and track the minimized $\mathbb{KL}$ divergence. This will provide an estimate of the capacity of $Gen$'s reconstruction of single cells without a critic's input. Then, set $m$ to be a value close to minimized $\mathbb{KL}$ divergence after $n$ epochs. This feature is already implemented in our package, with a default VAE-only training of 10 epochs. In practice, we found that values of $m$ roughly close to this minimized divergence performed well.
2. Another strategy for choosing $m$ can be a rough grid-search starting from large values of $m$ (which could be the upper bound of $\mathbb{KL}$ values) and gradually going to 0.

### Network Architecture

#### 3.1 Encoder Network

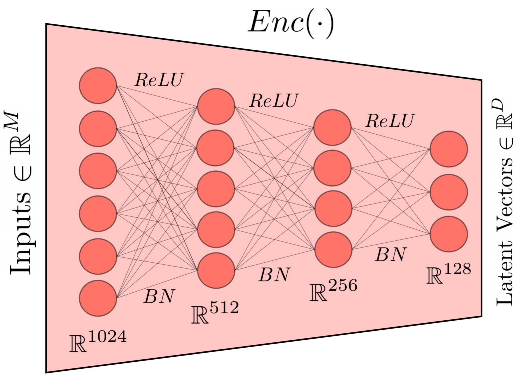


**Fig. S1.** The encoder network of ACTIVA.

$Enc$ consists of fully connected layers, with Rectified Linear Units $(ReLU)$ [Nair and Hinton (2010)] as the activation between two layer, where we also perform batch normalization operation in [Ioffe and Szegedy (2015)] (denoted as $BN$) after $ReLU$. The input to the network is $x \in \mathbb{R}^M$, which goes through the network with layers $\{1024, 512, 256, 128\}$, as shown in Fig. S1. Adam [Kingma and Ba (2015)] optimizer is used with a learning rate $lr = 0.0002$, and moving average decay rates $\beta_1 = 0.9$, $\beta_2 = 0.999$. Gradients are calculated on mini-batches of size 128, with the adversarial constant $m = 110$.

#### 3.2 Generator Network

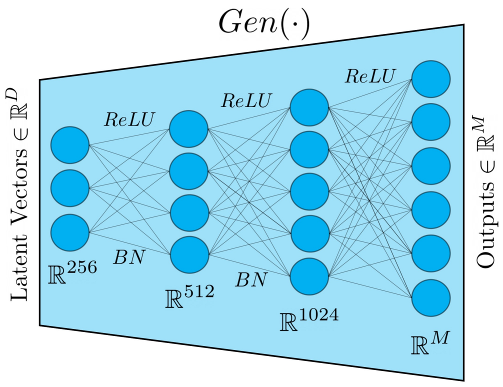


**Fig. S2.** Architecture of ACTIVA's generator network.

The generator network mirrors the encoder network, consisting of an input latent vector $z \in \mathbb{R}^D$ going through the layers $\{256, 512, 1024, M\}$ as shown in Fig. S2. Note that in the last layer of the generator (mapping from 1024 to $M$), we use

$ReLU$ without $BN$. This is because we want to ensure that all generated values are non-negative. Similar to the encoder, we optimize the network using Adam with a learning rate $lr = 0.0002$, $\beta_1 = 0.9$, $\beta_2 = 0.999$. Gradients are calculated on 128-cell mini-batches.

#### 3.3 Automated Cell Type Network (ACTINN)

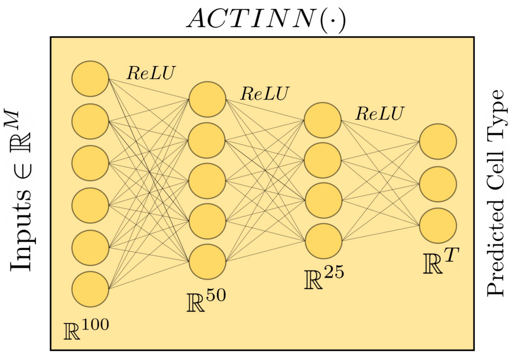


**Fig. S3.** Automatic cell-type identification network of ACTIVA, which is ACTINN.

Ma and Pellegrini (2019) use a fully connected neural network architecture for supervised classification of cell-types. scRNAseq data is usually high dimensional and often sparse, making neural network a promising method for analyzing such data. We implemented ACTINN in PyTorch, and used it for identifying cell-types and conditioning ACTIVA. Our implementation has same architecture as in Ma and Pellegrini (2019) which was implemented in TensorFlow (S3). Cross entropy is used to measure the loss between the predicted classes and the actual cell types. Optimization is done with Adam and an exponential "staircase" decay is used, with initial learning rate being $lr = 0.0001$ and a decay rate of 0.95 applied after every 1000 optimization steps. Our implementation of ACTINN in PyTorch differs from the original implementation in two ways: (1) we do not use a SoftMax layer between the last hidden layer and the output layer (due to the implementation of cross-entropy in PyTorch), and (2) we train for fewer number of epochs (between 5-10 as opposed to 50 epochs in the original implementation). Although we train for fewer epochs that Ma and Pellegrini (2019), we did not notice a drop in the accuracy of our model; that is, our results on 68K PBMC closely match the results found by Abdelaal *et al.* (2019) (results are shown in Tables S4-S5). Gradients are calculated on mini-batches of size 128.

### Downsampling and Data Augmentation

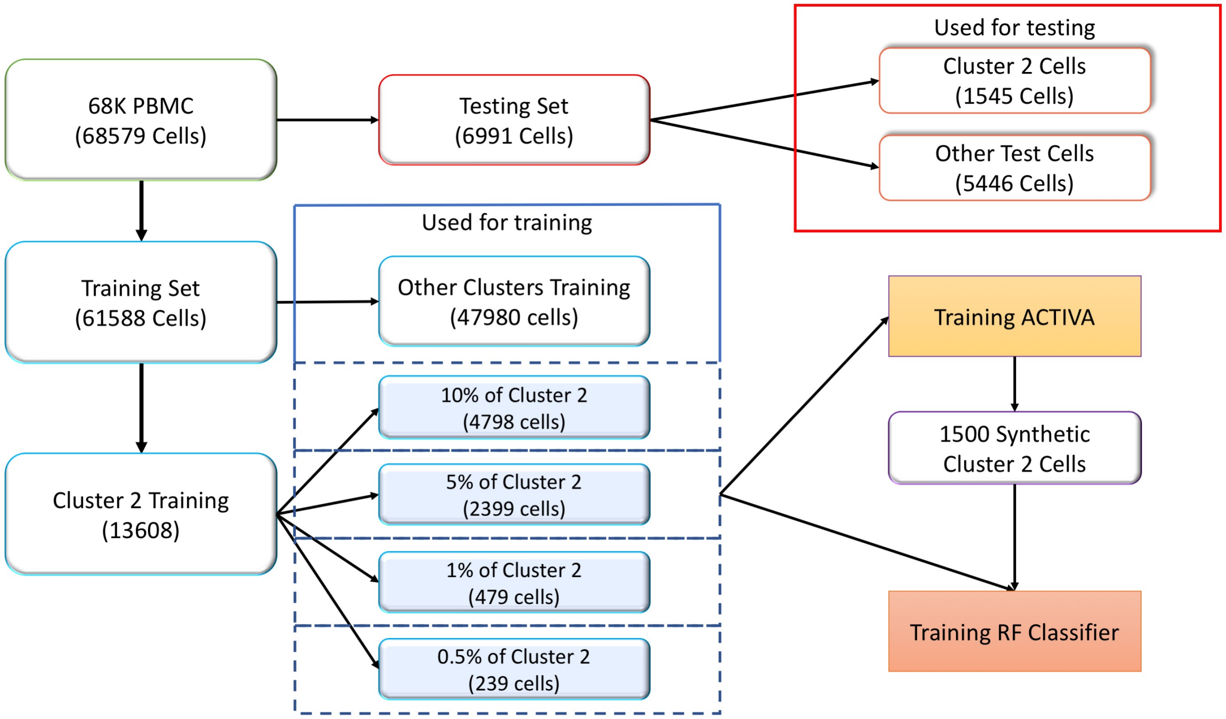


**Fig. S4.** Downsampling process for evaluating the impact of data augmentation with ACTIVA (Test splits shown in red frames and Training splits are in blue frames, with ACTIVA generated in purple frame). We first separate the test set from the training set, and subsequently label cluster 2 cells to differentiate them from all other clusters. Cells from all other clusters (besides cluster 2) are used in all training and testing modes. For cluster 2 cells, we randomly subsample a fraction of the cells (10%, 5%, 1% or 0.5%) and use this subset in addition to all other cells to train ACTIVA. In other words, ACTIVA and the RF will include (i) training data from all other clusters and (ii) one of the downsampled version of cluster 2 cells (highlighted in light blue). For the performance evaluation of RF without data augmentation ("no-augmentation"), we only use the desired cluster 2 subset and all other training cells to train the classifier. For training mode ACTIVA augmentation, we generate 1500 cells for data augmentation, and add to the training cells we used in "no-augmentation" mode. So for training the RF in augmentation mode, we use (i) training data from all other clusters, (ii) one of the downsampled version of cluster 2 cells (highlighted in light blue) and (iii) 1500 ACTIVA generated cells [trained on the same donwsampled data in (ii)].

### Additional Results

#### 5.1 Accuracy of the Classifier on Each Dataset

As mentioned in the main manuscript, the sub-population cell generation depends on correctly classifying the cell-types. Here, we show that ACTIVA's classifier, ACTINN, can classify rare-cell population, with training cells as few as 71 cells (see Table S5). We did observe that in some cases, if the number of training samples for a specific cluster is extremely low, then ACTIVA does not learn that cell-type (as it is expected from any machine learning model); for example, the classifier does not learn the 10th cell-types of the PBMC data, since there were only 19 training cells available (out of 61K training cells). However, for cluster 2 cell population, having 50 cells resulted in an F1 score of 0.74. This was useful for studying the impact of data augmentation with ACTIVA, as described in Section 5 of the main manuscript.

**Table S3.** Accuracy of ACTIVA's classifier network (ACTINN) on the test sets of Brain Small and 68K PBMC. Three metrics were used: (1) Accuracy-number of correct predictions over all predictions; (2) *F1 Score (Non-Weighted)*: unweighted mean of per-label accuracy (not counting for cell-type imbalance); and (3) *Weighted F1 Score*: per-type accuracy but weighted by the number of cells for each cell-type.

| Test Set | Accuracy | F1 Score (Non-Weighted) | Weighted F1 Score |
|---|---|---|---|
| Brain Small | 0.9649 | 0.9674 | 0.9654 |
| 68K PBMC | 0.9223 | 0.7448 | 0.9216 |

**Table S4.** Here we present the accuracy of ACTIVA's classifier network on the test and training set for Brain Small dataset (see main text Section 4.3 and Fig. 4).

| Cluster | Testing Cells | Test Precision | Test Recall | Test F1-Score | | Training Cells | Train Precision | Train Recall | Train F1-Score |
|---|---|---|---|---|---|---|---|---|---|
| 0 | 978 | 0.97 | 0.97 | 0.97 | | 8808 | 0.99 | 1.00 | 0.99 |
| 1 | 304 | 0.98 | 0.99 | 0.98 | | 2738 | 0.99 | 1.00 | 0.99 |
| 2 | 271 | 0.90 | 0.93 | 0.92 | | 2439 | 0.98 | 0.99 | 0.98 |
| 3 | 182 | 0.99 | 0.96 | 0.97 | | 1646 | 1.00 | 0.98 | 0.99 |
| 4 | 141 | 0.99 | 0.94 | 0.97 | | 1271 | 1.00 | 0.99 | 0.99 |
| 5 | 59 | 0.98 | 0.98 | 0.98 | | 535 | 1.00 | 0.99 | 1.00 |
| 6 | 35 | 1.00 | 0.94 | 0.97 | | 323 | 1.00 | 0.97 | 0.98 |
| 7 | 27 | 1.00 | 0.93 | 0.96 | | 243 | 0.99 | 0.99 | 0.99 |

**Table S5.** Here we present the accuracy of ACTIVA's classifier network on the test and training set for 68K PBMC dataset(see main text Section 4.3 and Fig. 4).

| Cluster | Testing Cells | Test Precision | Test Recall | Test F1-Score | | Training Cells | Train Precision | Train Recall | Train F1-Score |
|---|---|---|---|---|---|---|---|---|---|
| 0 | 1791 | 0.89 | 0.90 | 0.90 | | 15768 | 0.99 | 1.00 | 1.00 |
| 1 | 1545 | 0.88 | 0.87 | 0.88 | | 13608 | 1.00 | 0.99 | 1.00 |
| 2 | 1515 | 0.93 | 0.96 | 0.95 | | 13344 | 1.00 | 1.00 | 1.00 |
| 3 | 697 | 0.91 | 0.87 | 0.89 | | 6145 | 1.00 | 0.99 | 1.00 |
| 4 | 483 | 0.99 | 0.98 | 0.99 | | 4258 | 1.00 | 0.99 | 1.00 |
| 5 | 466 | 0.97 | 0.97 | 0.97 | | 4105 | 1.00 | 1.00 | 1.00 |
| 6 | 413 | 1.00 | 1.00 | 1.00 | | 3644 | 1.00 | 1.00 | 1.00 |
| 7 | 71 | 0.98 | 0.82 | 0.89 | | 626 | 1.00 | 0.95 | 0.97 |
| 8 | 8 | 0.00 | 0.00 | 0.00 | | 71 | 1.00 | 0.68 | 0.81 |
| 9 | 2 | 0.00 | 0.00 | 0.00 | | 19 | 0.00 | 0.00 | 0.00 |

### 5.2 Gene Expressions

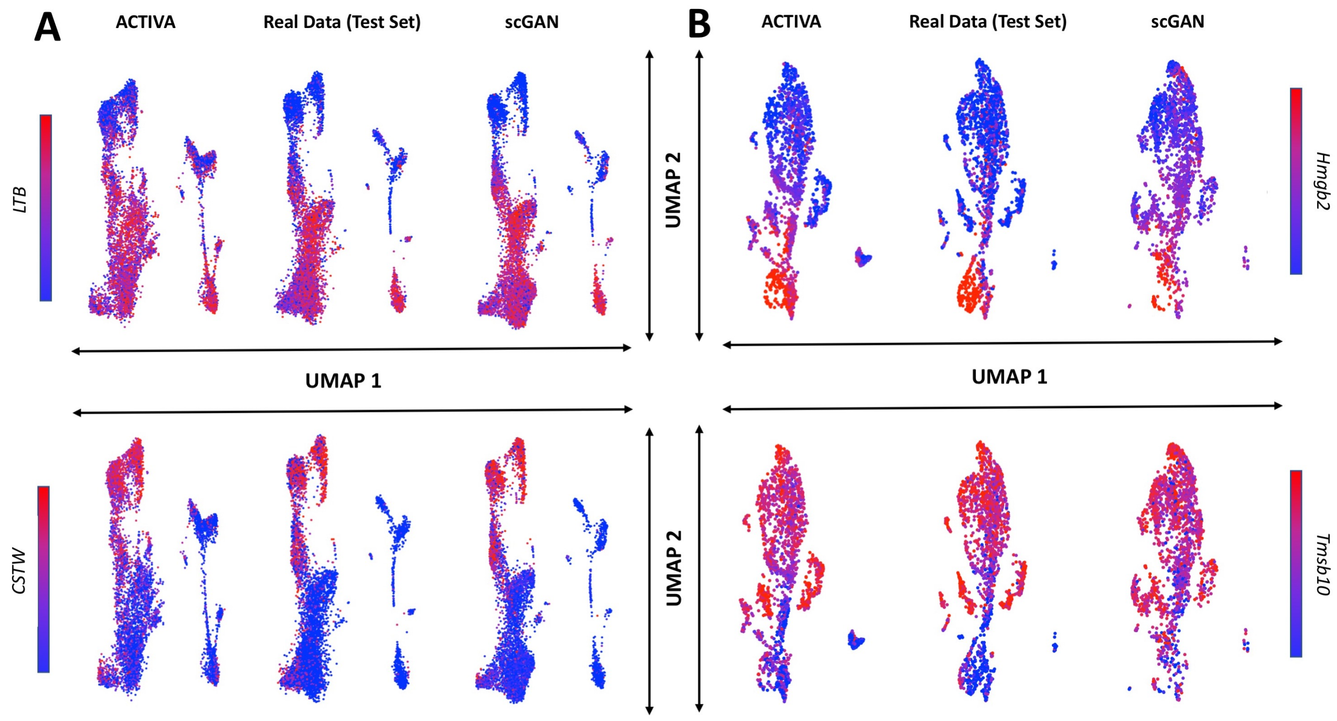


**Fig. S5.** UMAP of Synthetic Cells (generated by ACTIVA and scGAN with a subset of real data as training data) compared to real data (not used in training) colored by gene expression. Column (A) . **Column A**: here we present results from 68K PBMC data with a UMAP of 6991 cells generated by ACTIVA and scGAN, respectively, along with the test set, colored by the gene expression of two markers genes. **Column B**: we show the results on Brain Small test set and 1997 ACTIVA and scGAN generated cells, colored by the gene expression of two marker genes. For both datasets, we see that cells generated by ACTIVA resemble the real data well while more diversity is present among the generated cells.

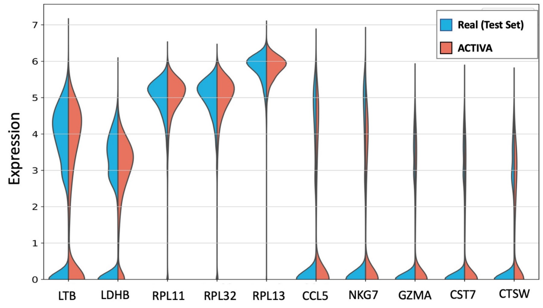


**Fig. S6. Logarithmic expression of the top marker genes**. First 5 are for cluster 1 and last 5 for cluster 2 in 68K PBMC test set (in blue) and ACTIVA generated cells (in red). The similarity between the distributions indicates that ACTIVA has learned the underlying marker gene expression.

#### 5.3 Manifold Analysis

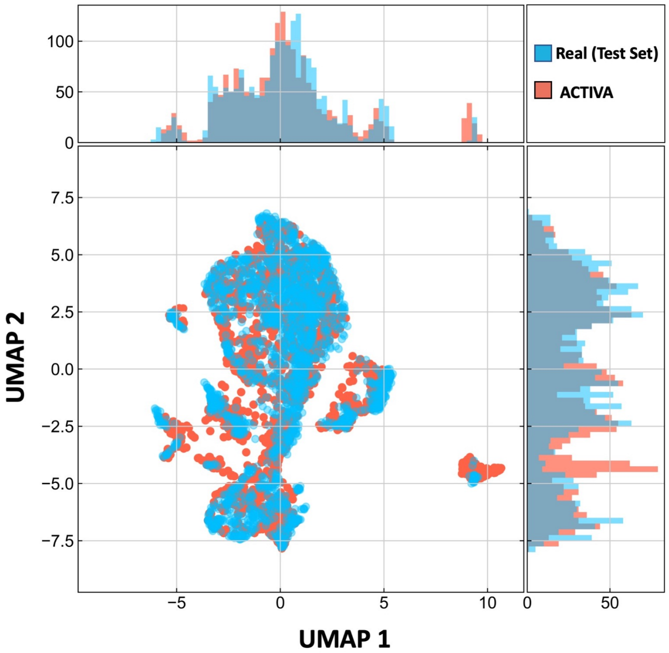


**Fig. S7.** UMAP of cells generated by ACTIVA compared to real test cells (not used in training) from 20K Brain Small. The histograms on top and right of the UMAP plot display the counts of cells on the horizontal and vertical axis, respectively. This figure shows that ACTIVA has learned the underlying distribution of the real data while having some diversity in the generated samples (which is desired for generative models).

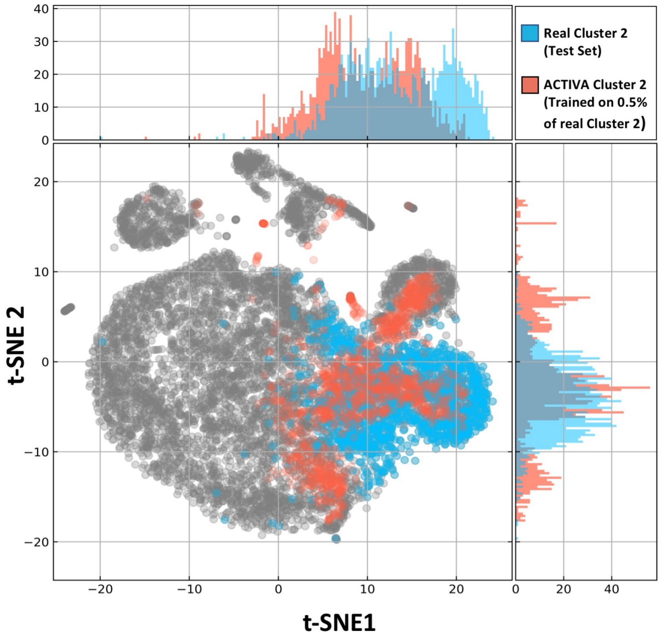


**Fig. S8.** t-SNE of all the cells in the test set (in grey), real cluster 2 cells (in blue), and ACTIVA generated cells when trained with only 0.5% of cluster 2 cells (in red). This figure illustrates that ACTIVA learns to generate a sub-population even when it is trained on 239 cells (out of 7915 training cells). This qualitative evaluation, combined with the results shown in the manuscript, show promising application of ACTIVA for improving downstream classification of rare populations.
